# Pose prediction accuracy in ligand docking to RNA

**DOI:** 10.1101/2023.05.22.541762

**Authors:** Rupesh Agarwal, T. Rajitha Rajeshwar, Jeremy C. Smith

## Abstract

Structure-based virtual high-throughput screening is used in early-stage drug discovery. Over the years, docking protocols and scoring functions for protein-ligand complexes have evolved to improve accuracy in the computation of binding strengths and poses. In the last decade, RNA has also emerged as a target class for new small molecule drugs. However, most ligand docking programs have been validated and tested for proteins and not RNA. Here, we test the docking power (pose prediction accuracy) of three state-of-the-art docking protocols on ∼173 RNA-small molecule crystal structures. The programs are AutoDock4 (AD4) and AutoDock Vina (Vina), which were designed for protein targets, and rDock, which was designed for both protein and nucleic acid targets. AD4 performed relatively poorly. For RNA targets for which a crystal structure of a bound ligand is available, and the goal is to identify new molecules for the same pocket, rDock performs slightly better than Vina. However, in the more common type of early-stage drug discovery setting, in which no structure of a ligand:target complex is known, rDock performed similar to Vina, with a low success rate of ∼27 %. Vina was found to bias for ligands with certain physicochemical properties whereas rDock performs similarly for all ligand properties. Thus, for projects where no ligand:protein structure already exists, Vina and rDock are both applicable. However, the relatively poor performance of all methods relative to protein target docking illustrates a need for further methods refinement.

## Introduction

The drug discovery process is expensive and time-consuming^1-3^. However, in the last few decades, computational resources have been used to expedite different steps of the process. One such step is in early-stage drug discovery, for which virtual high-throughput screening (v-HTS) with molecular docking has evolved into an essential tool in fast and cost-effective small molecule discovery and drug optimization. Molecular docking is often used to screen large chemical libraries for binding to a certain drug target. The method calculates the most favorable position (pose) of a ligand in a binding site and ranks the binding energies. Many novel bioactive molecules have been identified using this method^4^.

Historically, proteins have been considered as the primary targets of small molecule drugs. However, in the last two decades, since the Encyclopedia of DNA Elements (ENCODE) project^5^, RNAs have also also emerged as therapeutically relevant. 90% of the genome is transcribed into RNA, but the majority of this does not transcribe into proteins^6, 7^. These non-coding RNAs have been discovered to have diverse functions and play important roles in several diseases^8-10^. Recent work has shown that, similar to proteins, small molecules can be used to target RNA and that, perhaps unsurprisingly, similar computational methods can be used as well^11-18^.

There are many protein docking programs available, and each has its own pros and cons. AutoDock methods (AutoDock4 (AD4) ^19, 20^ and AutoDock Vina (Vina)^21^) are among the most popular and are open source and easy to use. Moreover, AD4 and Vina perform well in protein benchmarking studies^22^. However, like most existing docking protocols, AutoDock methods were designed for only protein-ligand docking. In contrast, another popular method, rDock^23^ was designed for both protein and nucleic acids and is one of the most popular and reliable docking methods for the latter. The precursor of rDock is the program RiboDock^24^, designed initially for virtual screening of RNA targets. The rDock scoring functions have been validated against both protein and RNA targets.

Several previous papers which have compared docking methods for RNA systems.^16, 25, 26^ Here, we ask the question, using a small sub-set of methods, as to in which circumstances RNA-oriented programs improve upon protein-oriented ones as concerns pose prediction in docking to RNA. We perform a detailed and systematic comparison of rDock, AD4 and Vina on a set of ∼173 RNA-ligand crystal structures, which is the largest dataset considered thus far for any comparison. We not only compare the default parameters of these methods but deep dive into testing different parameters and flavors of docking protocol within each method.

We performed two analyses: of the ‘docking power,’ *i*.*e*., the RMSD between the docked ligand pose and crystal ligand pose of the complex, and the “success rate” (*i*.*e*., the fraction of docked ligands within a given RMSD cut-off value from the reference ligand coordinates). We find that in cases where the docking box (i.e., search space) is not based on an existing ligand-bound crystal structure, the docking power of Vina was comparable to that of rDock. AD4 performed comparatively worse. In contrast, rDock performed significantly better than both AD4 and Vina when a ligand-bound crystal structure is known and used to define the docking search box, which can be the case in later hit expansion and lead optimization steps. Furthermore, unlike rDock, Vina performs better for only ligands within a certain range of physicochemical properties, including hydrogen bond acceptor/donor and molecular weight, which are considered important properties for drug discovery.

## Methodology

### Docking Dataset

The benchmark dataset consisted of 194 available crystal structures of RNA-ligand complexes derived from previous studies^27^. These were downloaded from the RCSB Protein Data Bank (PDB); the PDB IDs of all RNA−ligand complexes used are provided in **Table S1** of the supporting information. The PDB files were pre-processed to remove all cofactors and metal ions and any additional structures/chains present for all docking calculations. The docking site of each target was determined based on the position of the co-crystalized ligand within the active pocket. The PDB files of the receptors and ligands were converted into the required (PDBQT, SDF, MOL2) format using Babel and Mgltools. Due to the limitations of the conversion tools and docking programs, the results reported in this paper were computed for only 173 of the 194 RNA-ligand complexes for which docking was successfully completed for all docking programs.

### Docking parameters

#### AutoDock4 (AD4) and AutoDock Vina (Vina)

The center of the search box was set as the center of geometry of the crystallized ligand and the optimized size of the box (X) was calculated using eBoxsize^28, 29^ for both programs. Docking using Vina included box sizes of X and 2X with exhaustiveness kept at default (=8) and 25. For AD4, the AutoGrid program was employed in the construction of grid parameter files and atom-specific affinity maps, and the grid spacing was kept at 0.375 Å. The genetic algorithm (GA) run was selected as 10 with the population size and the number of generations 150 and 27000, respectively. AD4 docking was performed using box sizes of X and 2X. All other parameters for both the programs were kept at default.

#### rDock

The main components of the rDock scoring function are a van der Waals potential, an empirical term for attractive and repulsive polar interactions, and an optional desolvation potential that uses a weighted solvent accessible surface area. There are two methods to define the docking search space in rDock:

1. A two-sphere method is used to identify well-defined cavities in a receptor protein that are suitable for drug design. The method involves placing a grid over the cavity mapping region and using large spherical probes to eliminate flat and shallow regions, leaving behind well-defined cavities. Probes of smaller radii are then used to map the remaining unallocated grid points and identify the accessible regions. The final selection of cavity grid points is divided into distinct cavities that meet user-defined filters for minimum volume and maximum number of cavities.
2. The reference ligand method involves defining a docking volume around a known ligand’s binding mode by placing a grid over the cavity mapping region that encompasses overlapping spheres centered on each atom of the reference ligand. Grid points outside the overlapping spheres and within the receptor’s occupied volume are excluded, and small probes are placed on each remaining grid point to check for accessibility. The final selection of cavity grid points is divided into distinct cavities and filtered based on user-defined criteria such as the minimum volume and maximum number of cavities.

Docking poses were generated using the above two cavity defining methods implemented in rDock. The reference ligand method was used to define a docking volume of a given size around the binding mode of a known ligand, with sphere radii of 6 and 10 Å. One hundred poses were generated with each sphere radius. The two-sphere method, referred to as TS, was employed with outer sphere radii set to 6 and 10 Å. In addition to these, a radius of 4 Å was included in our calculations. Moreover, docking was performed with two scoring functions – the standard scoring function and the dock_solv (desolvation scoring function) available in rDock. In the desolvation scoring function, the repulsive polar term from the standard scoring function is replaced by a more finely parameterized desolvation potential based on a Weighted Solvent-Accessible Surface Area (WSAS) model. All other docking parameters were kept as default (See Supporting information).

Overall, we used six docking protocols: AD4, and Vina and four rDock protocols combining two cavity defining methods (Reference Ligand (RL) and Two Sphere (TS)) with two scoring functions (dock (standard scoring function) and dock_solv (desolvation scoring function)). Hereafter, the six docking protocols will be referred to as AD4, Vina, RL/dock, RL/dock_solv, TS/dock, TS/dock_solv.

#### Analyses

To evaluate the docking power of each program the RMSD (root mean square deviation) between the predicted (docked) pose and the native (crystal) binding pose of the ligand was calculated. The conformation with the highest docking score is referred to as the top scored pose (*top pose*) and the conformation that is closest to the native binding pose of the top five scored poses is referred as ‘*best of the top 5 poses’*. All RMSD values were calculated using the obrms module of Openbabel^30^. The success rates were evaluated for the docking programs with threshold RMSDs of 1.5 Å, 2.0 Å, and 2.5 Å relative to the crystal structure. A threshold RMSD of 2.5 Å was considered for evaluation of the performance/success rate of docking programs for RNA-ligand complexes as the ligand pockets in RNA tend to not be closed cavities and are less hydrophobic^25, 31-33^.

## Results and Discussion

### Docking power

Firstly, to compare the docking powers for all six docking methods we calculated the RMSD between the top pose or the best of top 5 poses and the crystal pose. The RMSD distributions are shown in **Figure 1**, and the median RMSD (mRMSD) values are reported in **Table 1-2**. It is observed that for the top poses Vina and AD4 show similar, bimodal RMSD distributions with peaks at both ∼1.5 Å and ∼7 Å and with a mRMSD of ∼ 4.9 Å. When the best of 5 poses is considered, the bimodal distribution is observed to be retained for AD4 but a unimodal distribution peaked ∼2 Å is observed for Vina, indicating that Vina performs better than AD4. The mRMSD of AD4 and Vina for the best of the top 5 poses are 3.6 Å and 2.7 Å, respectively.

**Table 1.** Median RMSD (mRMSD) for Top Pose.

| Docking Program | AD4 | Vina | RL/dock, | RL/docksolv | TS/dock | TS/docksolv |
| --- | --- | --- | --- | --- | --- | --- |
| mRMSD | 4.92 Å | 4.91 Å | 2.83 Å | 5.84 Å | 5.97 Å | 6.11 Å |

**Table 2.** Median RMSD (mRMSD) for Best of Top 5 Poses.

| Docking Program | AD4 | Vina | RL/dock | RL/docksolv | TS/dock | TS/docksolv |
| --- | --- | --- | --- | --- | --- | --- |
| mRMSD | 3.56 Å | 2.75 Å | 1.94 Å | 4.57 Å | 5.34 Å | 5.63 Å |

**Figure 1:**
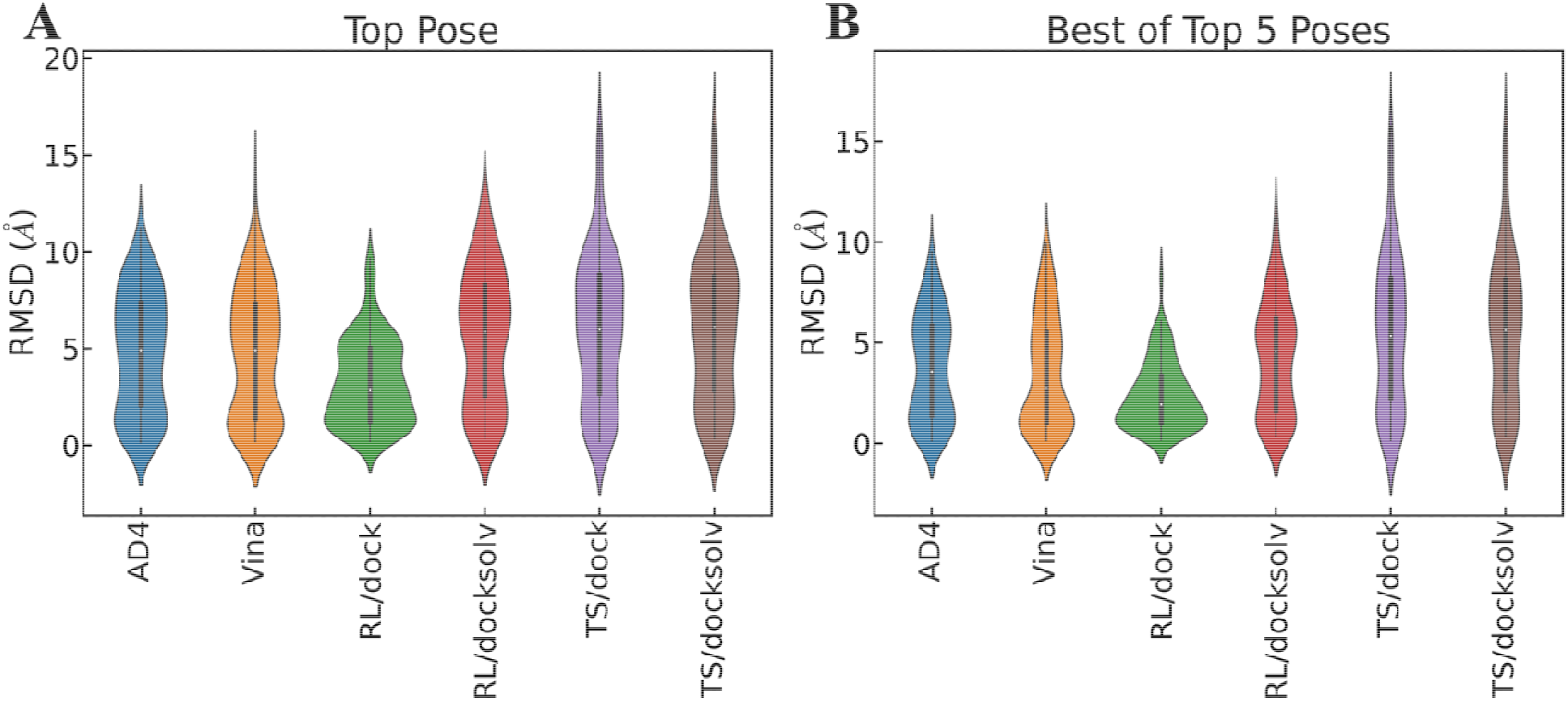
RMSD distributions for A) Top pose B) Best of Top 5 Poses are shown as violin plots for AutoDock family docking programs (AutoDock 4 and AutoDock Vina) and rDock docking program (Reference Ligand Method (RL) and Two Sphere Method (TS) with with dock and dock solv scoring functions).

Figure 1 also shows that for the top pose rDock RL/dock performs best with a comparatively higher peak around 2 Å and an mRMSD of 2.83 Å. The results of RL/dock are further improved upon consideration of the best of the top 5 poses with an mRMSD of 1.94 Å. RL/docksolv performs similar to the AutoDock methods, whereas poor performance is observed for TS/dock and TS/docksolv relative to the other methods, both for the top pose and the best of the top 5 poses. Comparison of the top pose RMSD distributions with the best of top 5 RMSD distributions further confirms that taking the best of the top 5 leads to better agreement with the experiment.

### Success rate

Next, we calculated the “success rate” of each of the methods, which we define as is the percentage of complexes for which the docking pose has an RMSD less than a given cutoff, of 1.5 Å, 2.0 Å, or 2.5 Å. The success rates of the docking programs for the top scored and best of top 5 poses for three RMSD thresholds are illustrated in **Figure 2**. The success rate for the top scored poses is less than or equal to 30 %, 35 % and 45 % for RMSD thresholds of 1.5 Å, 2.0 Å, and 2.5 Å, respectively, whereas the success rate for the best of top 5 scored poses is less than or equal to 40 %, 50 % and 65 % for RMSD thresholds of 1.5 Å, 2.0 Å, and 2.5 Å, respectively.

**Figure 2:**
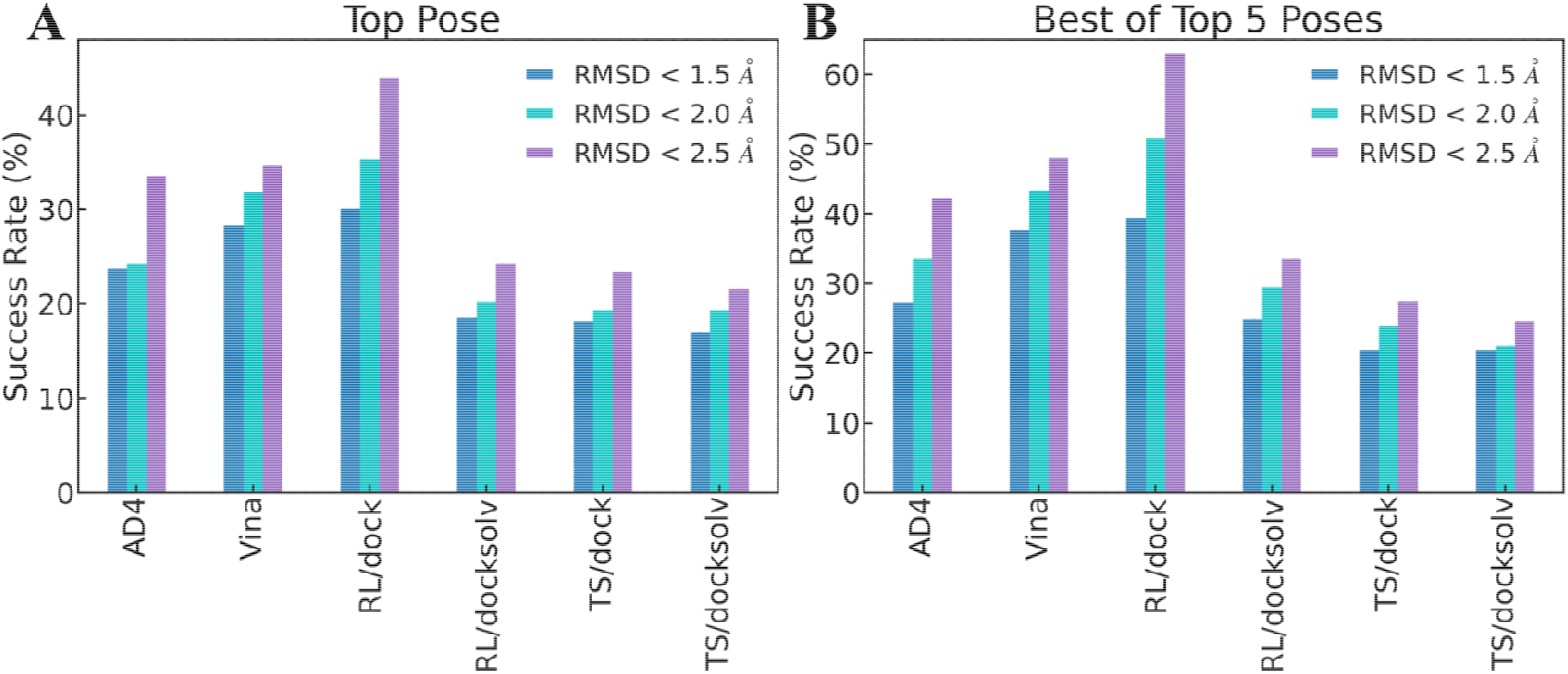
Success rates for A) Top pose B) Best of Top 5 Poses for AutoDock family docking programs (AutoDock 4 and AutoDock Vina) and rDock docking program (Reference Ligand Method (RL) and Two Sphere Method (TS) with with dock and dock solv scoring functions).

Though at the RMSD threshold of 1.5 Å none of the docking programs exceed 30 % success rate for the top pose or 40 % for the best of top 5 poses, it is interesting to note that the performance of Vina is comparable to rDock RL/dock. Also, for a 2.0 Å cutoff, the performance of Vina is quite comparable to that of rDock RL/dock, with ∼ 32% and ∼35 % for the top pose Vina and RL/dock, respectively, and ∼ 43% and ∼51 % for the best of 5. When increasing the RMSD threshold from 1.5 Å to 2.5 Å the success rate for the top pose of AD4 increases ∼15 % and becomes similar to that of Vina, at 2.5 Å RMSD. At the 2.5 Å RMSD threshold the top and best of 5 success rates of RL/dock are 44 % and 63 %, much higher than the other methods. Relatively low success rates are found for all three RMSD thresholds when using rDock RL/docksolv, TS/dock and TS/docksolv. Further, there is no significant improvement in the success rates of these three methods when the best of the top 5 poses is considered.

For any given RMSD threshold, the performance follows the following order: RL/dock > Vina > AD4 > RL/docksolv > TS/dock > TS/docksolv. Overall, RL/dock exhibits the best performance on ligand pose prediction for RNA targets but Vina’s performance is quite comparable to it. This indicates that Vina can be used as an alternative program for the virtual screening of RNA-ligand complexes. It is noteworthy, however, that the success rates observed here for RNA docking are only about half as good as are found for protein docking using the same protocols, indicating that there is a need to improve scoring functions for RNA docking^34-36^.

### Representative/Test cases

We then looked at RNA:ligand docked poses and overlaid them with their corresponding crystal poses for those protocols which show the best success rates i.e., rDock RL/dock, Vina, and AD4. **Figure 3** illustrates three different receptor-ligand pairs to highlight cases where one docking program performs significantly better than the other two and a case where all three docking programs perform similarly.

**Figure 3:**
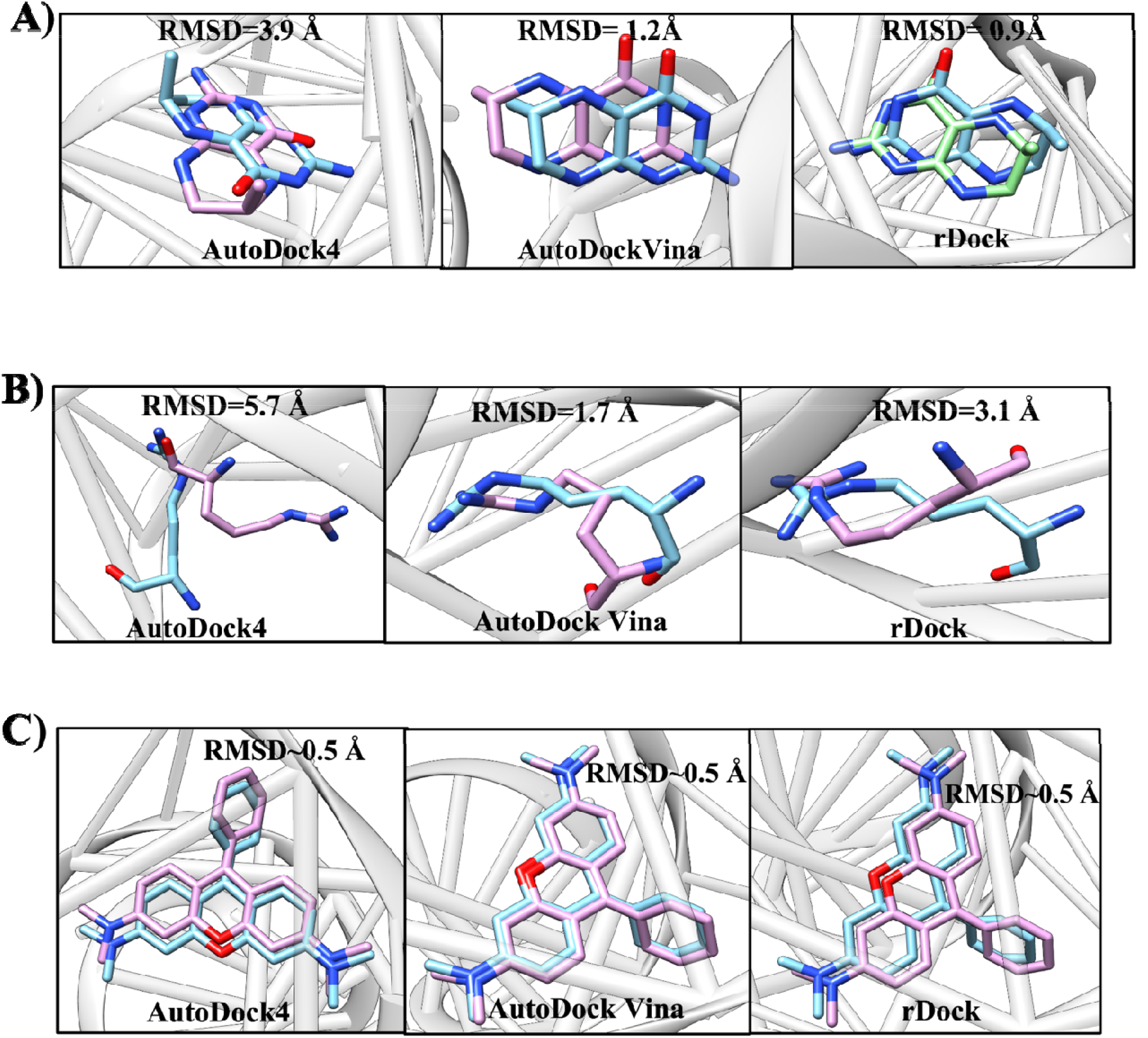
Top docking poses using A) 3SUX; B) 1AJU and C) 1F1T for docking programs AD4, AutoDock Vina, rDock (dock). RNA molecules are presented as a gray cartoon, ligands as sticks; crystal structure—cyan, solution found by docking program—pink. Heteroatoms are colored: O—red, N—blue.

**Figure 3A** shows a representative docking result for which rDock RL/dock performs best, PDB 3SUX, with the docked top pose of the ligand ∼0.9 Å from the crystal structure. Vina performs better than AD4: the overall crystal ligand conformation is obtained with Vina but the docked ligand pose is translated slightly, leading to an RMSD of ∼1.2 Å. The AD4 ring orientation does not match the crystal pose. **Figure 3B** illustrates the docking result for PDB 1AJU where the ligand is very flexible. We observe that Vina predicted a top pose closest to the crystal pose, with an RMSD∼1.7 Å. In this case, all docking programs got the pose wrong, but Vina and rDock RL/dock placed the terminal groups of the ligand in the correct pocket, whereas the pose obtained using AD4 flips the ligand completely. **Figure 3C** depicts docking results where all three docking programs predict the same top pose of the ligand, similar to the crystal pose, with an RMSD ∼0.5 Å.

### Ligand physicochemical bias in docking

To examine whether there is a dependence of the docking performance of any given program on the chemical properties of the ligands, analysis was made of physicochemical descriptors (including the molecular weight, number of heavy atoms, number of rotatable, amide, aliphatic bonds, number of H-bond donors and acceptors, MlogP (Moriguchin logarithm of the octanol/water partition coefficient), and polar surface area). This is in part because ligand physicochemical properties like molecular weight^37^ can have a significant influence on the performance of a docking program. For example, it was previously observed that Vina performs better than AutoDock4 in ranking large molecules, whereas they perform roughly equally good for ranking small molecules^38^.

Figure 4 shows representative distributions of the descriptors in three sets: i) ligands present in all the PDBs tested and ligands which performed best with, ii) rDock RL/dock, and iii) Vina. We chose these two as the success rates observed for RL/dock and Vina are high relative to other methods. All other distributions of descriptors are shown in **Figure S1**. The physicochemical descriptors for all the ligands of the dataset under consideration span a wide chemical space. rDock RL/dock performs relatively well with almost all the chemical space under consideration. However, the chemical space of ligands that perform well with Vina is found to be limited and is clearly distinguishable from that of rDock RL/dock. The ligands performing best with Vina have MolLogp ranging between 5 and -8, less than 10 rotatable bonds, less than 10 hydrogen bond acceptors/donors, a number of amide bonds less than 0.5 and a polar surface area less than 200 Å^2^. This analysis suggests that the Vina scoring function performs better for docking ligands within a specific domain of chemical properties and that it would therefore probably require tweaking of the scoring function terms to make it perform more broadly for RNA docking, whereas rDock, being already trained for RNA docking does not have an intrinsic bias for certain physicochemical features of RNA binding small molecules.

**Figure 4:**
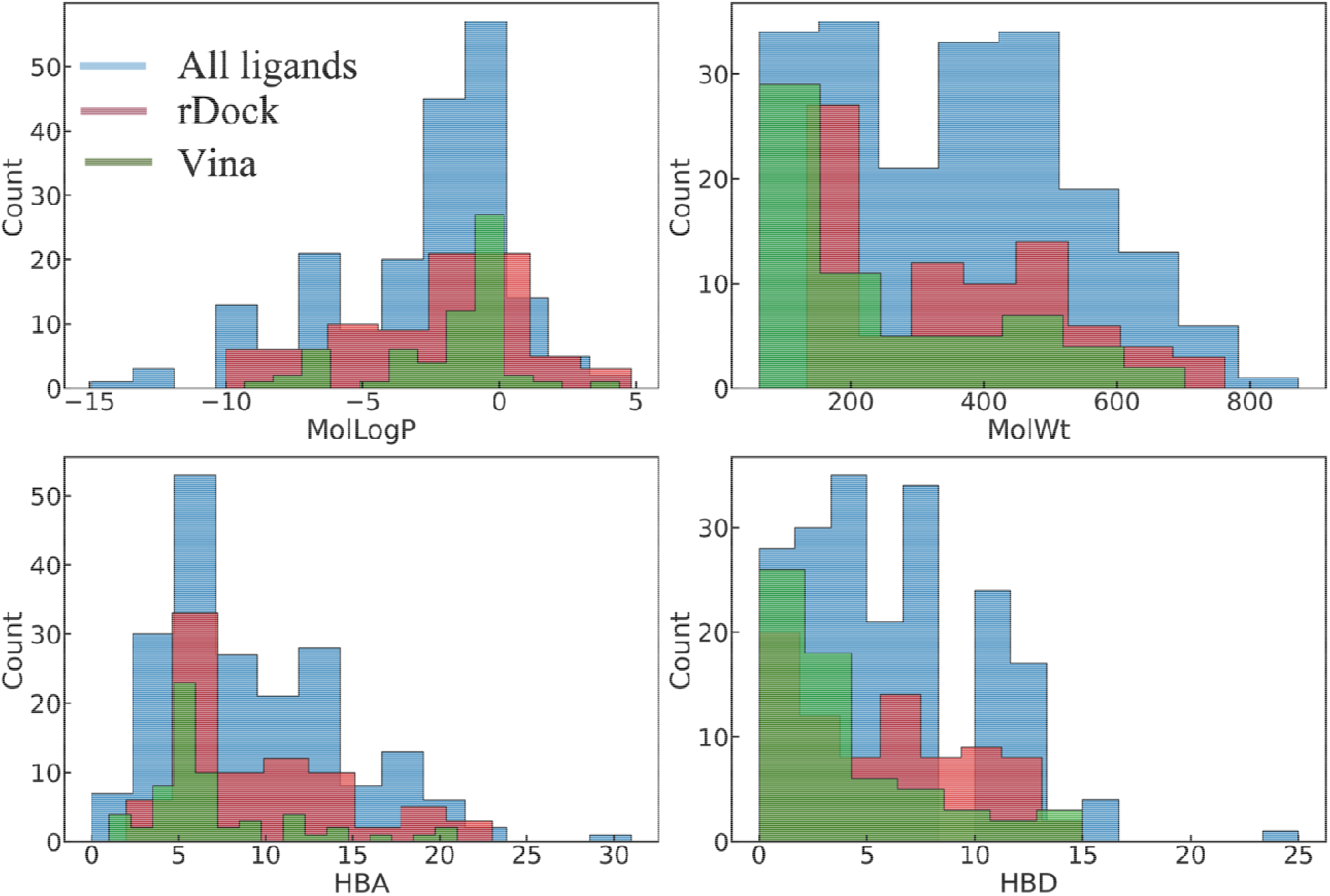
Representative physicochemical descriptors of the ligands that performed best with AutoDock Vina and rDock RL/dock docking programs.

### Tuning docking parameters of the docking tools

We also investigated whether the docking power of the tools can be improved by tuning docking parameters. For rDock, we changed the RADIUS parameter, which controls the cavity size included within the search space; for Vina, the box size and exhaustiveness were tuned, and for AD4 the box size was increased. In the case of the AutoDock methods the box size refers to the size of the docking search space and the exhaustiveness is the comprehensiveness of the search.

For rDock, we performed docking with two different RADIUS values - 4 Å and 6 Å – and compared the results with the manual default RADIUS value of 10 Å, for both the RL and TS methods with dock scoring, as, with the default values shown above this performed better than docksolv. The results, shown in **Figures 5 and S2** and tabulated in **Table S2**, show that mRMSD for RL/dock with RADIUS 4 Å and 10 Å performs better than 6 Å for both the top pose and for best of the top 5 poses. Similarly, there is no significant improvement observed for different success rates for RADIUS 4 Å and 10 Å, when compared to RL/dock with RADIUS 6 Å. This indicates that rDock RL/dock performs similarly with RADIUS 4 Å and 10 Å and much better than RADIUS 6 Å. Moreover, both the mRMSD (> 5 Å) and success rate (< 30 %) of the TS method did not change considerably upon varying the RADIUS parameter and with increasing RMSD threshold. Overall, the rDock RL method performs better than TS at all RADIUS values.

**Figure 5:**
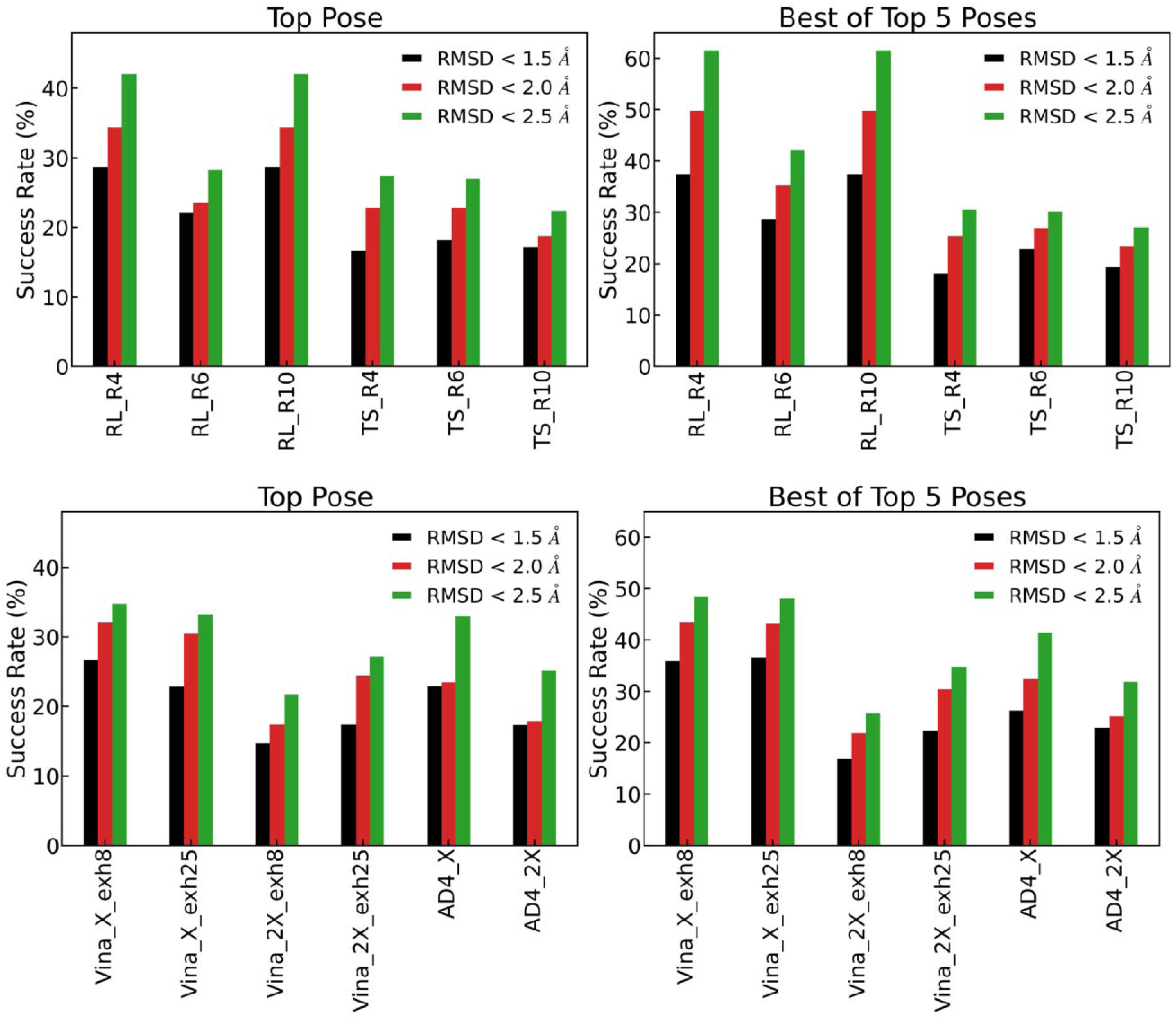
Top: Success rates for Top pose and Best of Top 5 Poses for AutoDock family docking programs (AutoDock 4 and AutoDock Vina) with exhaustiveness of 8 and 25 along with box sizes X and 2X. Bottom:Success rates for Top pose and Best ot Top 5 poses with rDock Reference Ligand Method (RL) and Two Sphere Method (TS) with dock scoring function and RADIUS 4 Å, 6 Å and 10 Å.

Similar to the RL method, the default box size in AutoDock tools was defined using the eBoxsize tool^28, 29^ which aims to calculate the optimal docking box size to maximize the accuracy of binding pose prediction by generating search boxes that are only big enough to include the crystallographic ligand pocket. For Vina and AD4, we doubled the box size and for Vina also considered two exhaustiveness values - 8 (default) and 25. The double sized search box is more realistic as the size of the box is often much bigger than the ligand in real-life drug discovery projects. We also changed the exhaustiveness value in the case of Vina to make the docking more comprehensive. The performance of the AutoDock tools decreased with increasing box size, as seen in **Figures 5 and S3** and **Table S3**, which is unsurprising as increasing the box size increases the available search space. However, increasing the exhaustiveness parameter from 8 to 25 in the case of Vina did not lead to any improvement (**Figure 5** and **Table S3**) thus showing that the limitation of Vina docking is not the exhaustiveness of the search but probably is more intrinsic, such as the scoring function or search algorithm. Overall, it seems that for a real-life structure-based drug discovery project to identify novel molecules i.e, without a crystal structure of a bound ligand, the performance of Vina is similar to that of rDock.

## Conclusions

RNAs have become important drug targets, and there has been an interest in understanding the limitations of conventional tools and developing new tools in using structure-based drug discovery methods to target RNAs. In this study, using an extensive benchmark dataset with 173 RNA-ligand complexes, the docking power (pose accuracy) of protein-based (AD4 and Vina) and protein-nucleic acid (rDock) docking programs were compared. rDock with dock and dock solv scoring functions perform the best overall in identifying the correct ligand binding poses if using the reference ligand (RL) method in which a crystal structure of a bound ligand is given as an input to define the search box. This result illustrates a drawback of rDock because the RL method requires coordinates of an a priori bound ligand in the receptor as an input to define the docking search space and can therefore only be applied for a well-defined pocket (*e*.*g*., with a ligand-bound crystal structure), which is mostly not available in virtual screening approaches. In contrast, the TS method has no such requirement and is thus more pragmatic and appropriate for identifying an initial hit molecule from screening. However, RL might therefore be useful for finding alternative molecules to an initial hit, i.e., as a strategy for finding multiple hits or hit expansion.

For the more common setting in which a ligand-bound crystal structure is not available AutoDock Vina performed similarly to rDock on pose prediction. However, the overall performance of both these tools is poor compared to protein-ligand docking. RNA docking is still an evolving field and has not yet been able to achieve accuracy similar to protein docking tools. Furthermore, we also found that Vina has an intrinsic bias towards certain ligand physicochemical descriptors. This relatively poor overall performance is not surprising since RNA is a negatively charged polymer and ligand binding often requires the participation of metal ions such as Mg^2+^. Scoring functions may need to reflect this. Also, ligand binding sites on RNA can be less deep and more polar, solvated, and conformationally flexible than for proteins, suggesting a possible need for modification of search methods.

## Supporting information

All Supplementary figures and tables

## Author Contributions

RA and RRT contributed equally to this work.

## Acknowledgment

This research used resources of the Compute and Data Environment for Science (CADES) at the Oak Ridge National Laboratory, which is supported by the Office of Science of the U.S. Department of Energy under Contract No. DE-AC05-00OR22725.

