## Supplementary material for "Pose prediction accuracy in ligand docking to RNA": All Supplementary figures and tables

**RNA-ligand complexes PDB IDs**

1AJU	1AKX	1AM0	1EI2	1EVV	1F1T	1FMN	1FUF	1FYP	1J7T	1LC4	1LVJ	1MWL	1NEM	1NTB	1O15	1O9M	1PBR	1Q8N	1QD3	1TN1	1TN2	1TOB	1U8D	1UTS	1UUD	1UUI	1XPF	1Y26	1YRJ	1ZZ5	2AU4	2B57	2BE0	2BEE	2CKY	2EES	2EET	2EEU	2EEV	2EEW	2ESJ	2ET3	2ET4	2ET5	2ET8	2F4S	2F4T	2F4U	2FCX	2FCY	2FCZ	2FD0	2G5K	2G5Q	2GDI	2GIS	2HO6	2HO7	2HOJ	2HOM	2HOO	2JUK	2L1V	2L94	2LWK	2M4Q	2MIY	2MXS	2N0J	2O3V	2O3W	2O3X	2O3Y	2OE5	2OE8	2QWY	2XNW	2YDH	3B4A	3B4B	3B4C	3C44	3D0U	3D2G	3D2V	3D2X	3DIG	3DIL	3DIM	3DIX	3DIY	3DIZ	3DJ0	3DJ2	3DVV	3E5E	3F2Q	3F2T	3FU2	3GCA	3GX2	3GX3	3GX5	3GX6	3GX7	3IQN	3IQR	3LA5	3NPN	3NPQ	3Q50	3RKF	3SD3	3SKI	3SKL	3SLM	3SLQ	3SUH	3SUX	3TD1	3TZR	4AOB	4E8N	4E8Q	4ERJ	4F8U	4F8V	4FAW	4FE5	4FEJ	4FEL	4FEN	4FEO	4FEP	4GPW	4GPX	4GPY	4JF2	4JIY	4KQY	4L81	4LVV	4LVW	4LVX	4LVY	4LVZ	4LW0	4LX5	4LX6	4NYA	4NYD	4P20	4P3S	4P5J	4P95	4PDQ	4QLM	4QLN	4RZD	4TS2	4TZX	4TZY	4XNR	4XW7	4XWF	4Y1I	4YAZ	4YB0	5BTP	5BWS	5C7W	5KPY

**rDock Parameters**

RECEPTOR_FLEX 3.0

SITE_MAPPER RbtLigandSiteMapper/ RbtSphereSiteMapper

RADIUS 4.0/6.0/10.0

SMALL_SPHERE 1.0

MIN_VOLUME 100

MAX_CAVITIES 1

VOL_INCR 0.0

GRIDSTEP 0.5

SCORING_FUNCTION RbtCavityGridSF

WEIGHT 1.0

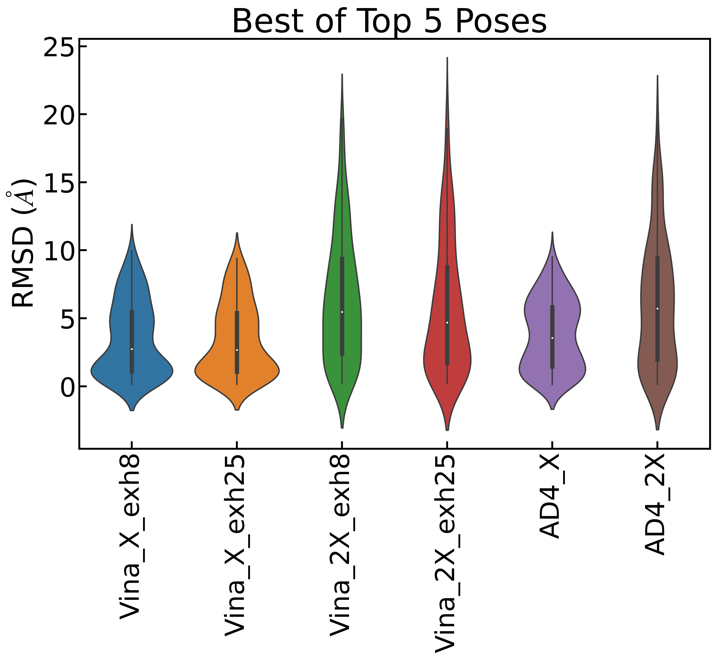

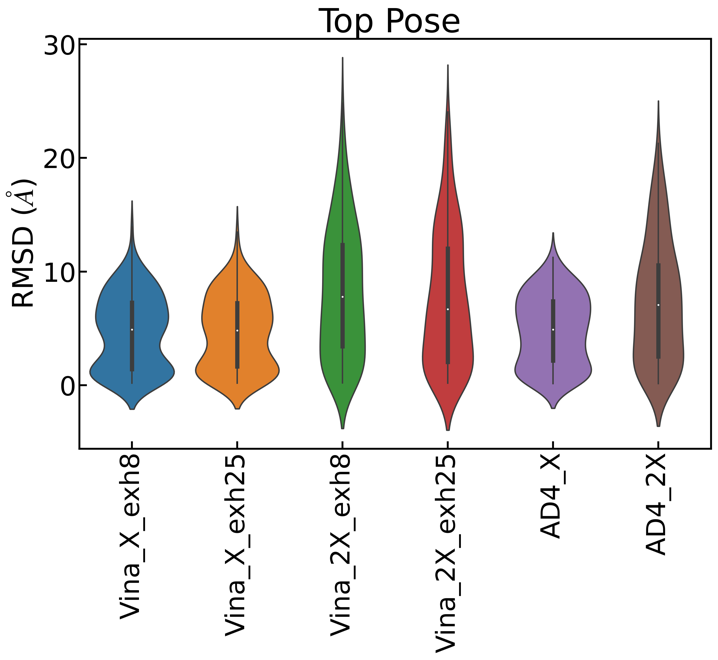

Figure S1: RMSD distributions for A) Top pose B) Best of Top 5 Poses are shown as violin plots for AutoDock family docking programs (AutoDock 4 and AutoDock Vina)

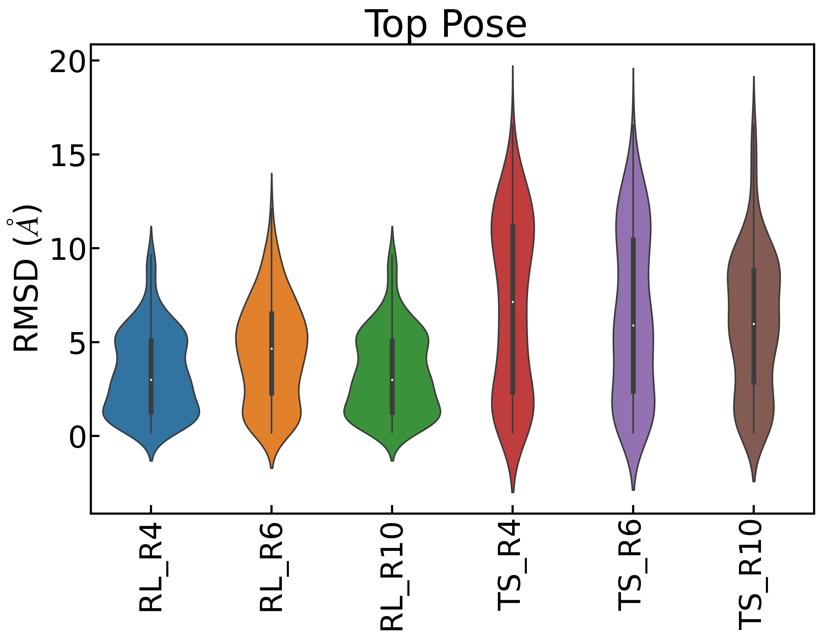

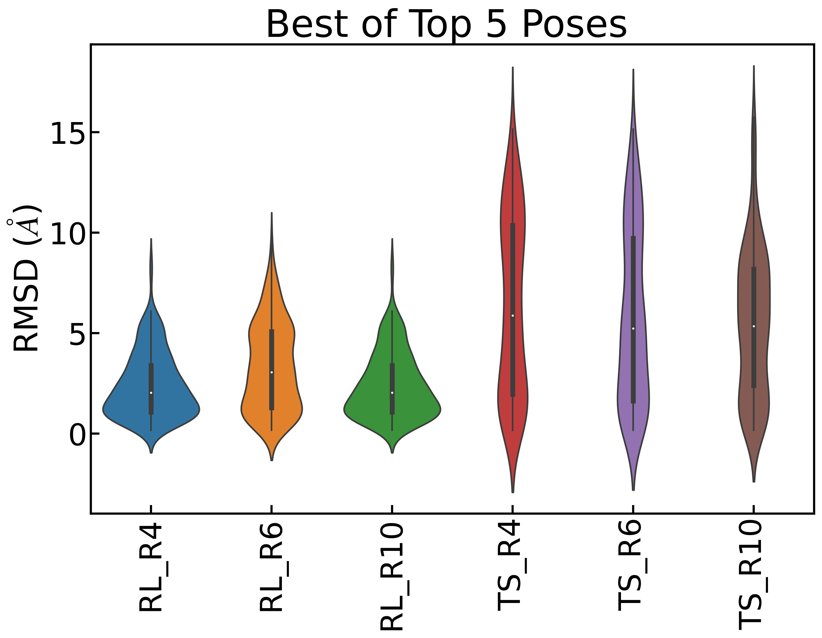

Figure S2: RMSD distributions for A) Top pose B) Best of Top 5 Poses are shown as violin plots for rDock docking program

**Table S1**

Top pose:

| rDock RL/dock | RL_R4 | RL_R6 | RL_R10 | TS_R4 | TS_R6 | TS_R10 |
| --- | --- | --- | --- | --- | --- | --- |
| mRMSD | 3.00 | 4.65 | 3.00 | 7.15 | 5.89 | 5.97 |

Best of Top 5 poses:

| rDock RL/dock | RL_R4 | RL_R6 | RL_R10 | TS_R4 | TS_R6 | TS_R10 |
| --- | --- | --- | --- | --- | --- | --- |
| mRMSD | 2.03 | 3.04 | 2.03 | 5.87 | 5.23 | 5.34 |

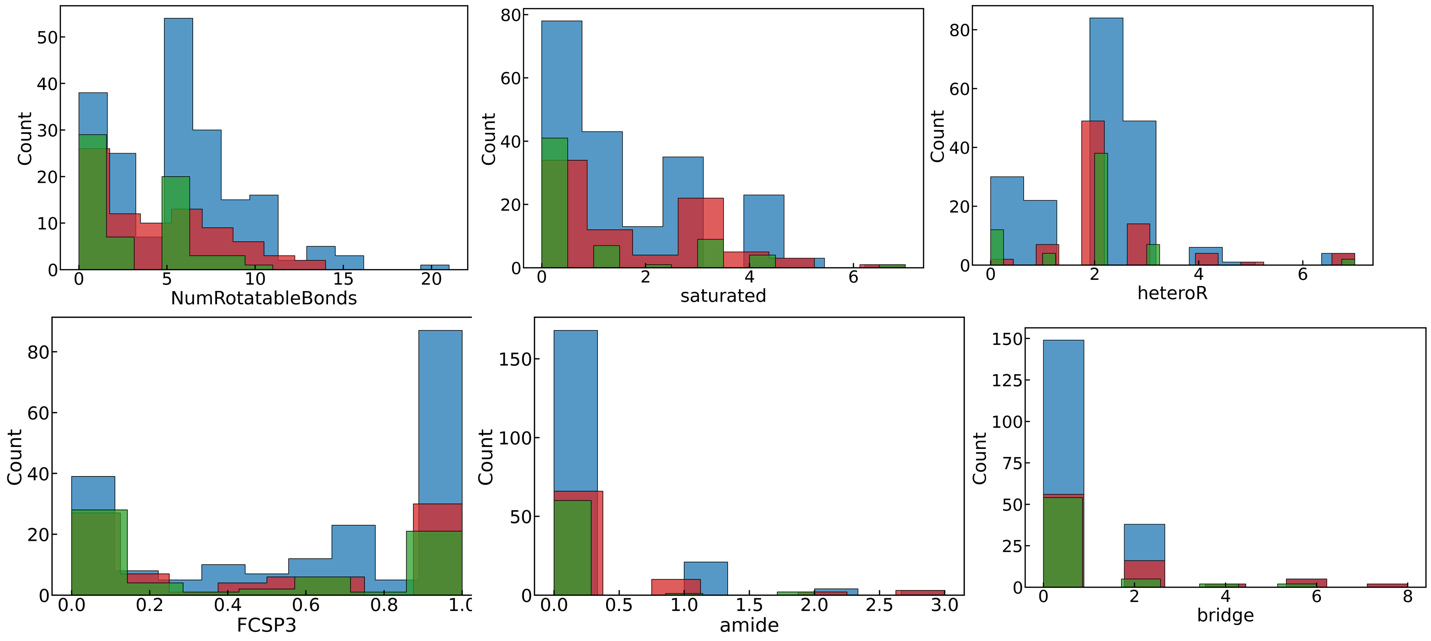

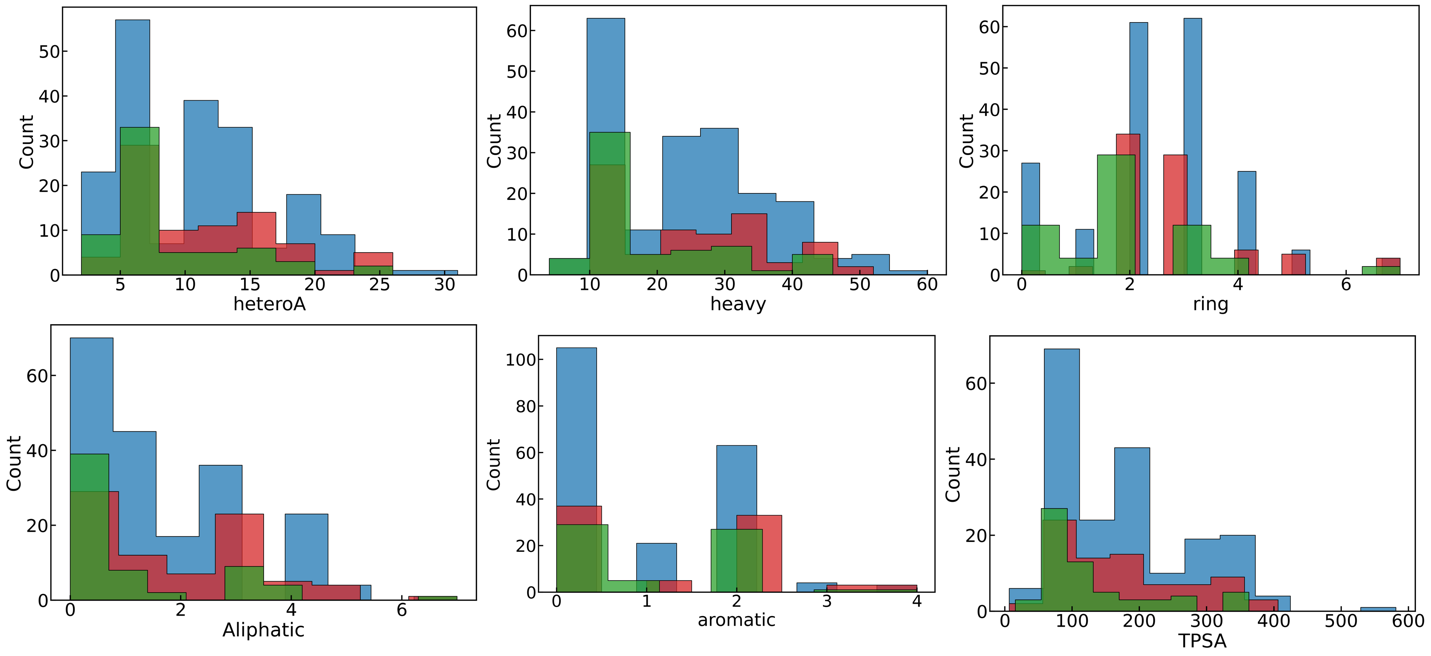

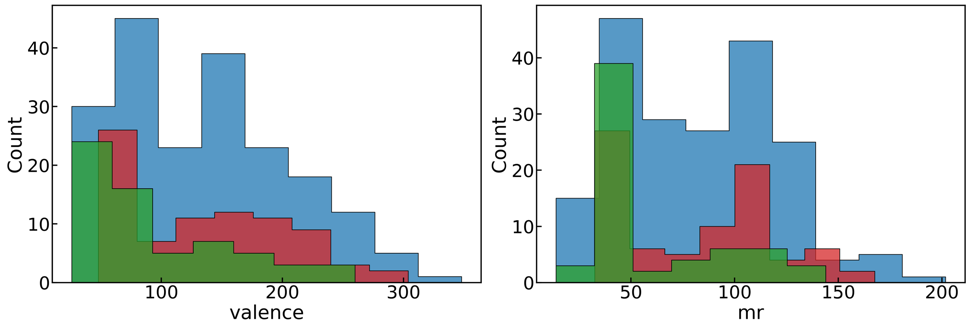

Figure S3: Different physicochemical descriptors of the ligands that performed best with AutoDock Vina and rDock RL/dock docking programs.
